# Land use land cover dynamics through time and their proximate drivers of change in a tropical mountain system: a case study in a highland landscape of northern Ecuador

**DOI:** 10.1101/2021.11.05.467450

**Authors:** Paulina Guarderas, Franz Smith, Marc Dufrene

## Abstract

Tropical mountain ecosystems are threatened by land use pressures, reducing the capacity of ecosystems to provide a large diversity of benefits to people and to be able to achieve them in the long term. The analysis of land use pressures is often superficial and very general, although they are characterized by numerous interactions and strong differences in their local dynamics. We used a variety of freely available geospatial and temporal data and methods to assess and explain patterns of land use land cover (LULC) change, focusing on native ecosystem dynamics, in a sensitive region of the northern Ecuadorian Andes. Our results demonstrate a dynamic and clear geographical pattern of distinct LULC transitions through time, explained by different combination of socio-economic factors, pressure variables and environmental parameters, from which ecological context variables, such as slope and elevation, were the main drivers of change in this landscape. We found that deforestation of remnant native forest and agricultural expansion still occur in higher elevations located, while land conversion toward anthropic environments were observed in lower elevations to the east of the studied territory. Our findings also reveal an unexpected stability trend of paramo and a successional recovery of previous agricultural land to the west and center of the territory which could be explained by agricultural land abandonment. However, the very low probability of persistence of montane forests in most of the studied landscape, highlights the risk that the remnant montane forests will be permanently lost in a few years, posing a greater threat to the already vulnerable biodiversity and limiting the capacity ecosystem service provisioning. The dynamic patterns through space and time and their explanatory drivers, found in our study, could help improve sustainably resource land management in vulnerable landscapes such as the tropical Andes in northern Ecuador.

## Introduction

Tropical mountain systems supply vital ecosystem services to millions of upland and lowland inhabitants [1,2], but are increasingly being transformed by human activities [3,4]. Tropical Andean mountain ecosystems are global hotspots of tropical biodiversity and habitat refugia [4,5] and are threatened by agricultural expansion and intensification, urbanization, pollution, mining, among other human activities, reducing the capacity of these ecosystems to provide a large diversity of benefits to people and guarantee their long-term sustainability.

Although the millennial human presence in the Andean mountain systems has impacted the history of landscape patterns in this region, mainly shaped by intensive traditional agriculture, which has been practiced for centuries [6], recent land-change patterns have been documented in this region [7–10], demonstrating varying patterns of landscape dynamism and heterogeneity [11]. Deforestation and agricultural intensification are the dominant transitions in many Andean systems [8,9], but forest recovery due to agricultural de-intensification and transitions between crops, pastures, and secondary vegetation have also been observed in these systems, yet in-depth multi-temporal change approaches are required to better understand this complexity in order to balance biodiversity conservation with human needs [7,9,12].

Distinct land use and land cover conversion (LULCC) patterns observed in the tropical Andes vary along demographic, socio-economic, cultural and technological factors that interact with biophysical features like elevation, topography, soils and climate parameters [3,13–15], operating across spatial, temporal, and organizational scales [13,14,16]. For example, increasing global demand for food and non-food crops can drive agriculture expansion in more fertile and flat land [7,17–19], where natural ecosystem recovery has been observed in abandoned marginal agricultural land [3,19,20].

Despite the documented useful insights on how different drivers can influence LULCC on specific landscape mosaics in tropical mountain systems, evidence from synthetical studies suggests that no universal link between cause and effect exists to explain LULCC, especially deforestation in tropical systems, where different combinations of various proximate causes and underlying driving forces in varying geographical and historical contexts could affect landscape changes [17,21].

Part of the key to understanding future changes in tropical mountain systems may come from better understanding LULCC pattern-dynamics across environmental gradients and along different temporal scales, in addition to deciphering the interactive effects of distinct anthropogenic influences on these kinds of processes [3], which could be exacerbated, given the high vulnerability to climate change of highland landscapes like the Ecuadorian Andes [22]. Understanding this complexity is fundamental for the development of policies and measures for landscape planning and management in highland ecosystems, where biodiversity conservation, sustainable use of natural resources and the supply of essential ecosystem services (eg. water or food) should be assured [23,24] not only for local inhabitants but also for downstream populations[19].

This study is unique in that it uses a variety of geospatial and temporal data and methods to assess and explain patterns of LULCC in a sensitive region of the northeastern Ecuadorian Andes, which comprises a landscape with distinct climatic conditions and management regimes along its elevation gradient, where floriculture crops and urban centres are emerging in an agricultural matrix, posing more pressure to remnant native ecosystems and their services. The objective of this paper was to assess the role of different drivers on LULCC in a highland landscape of Northern Ecuador from 1990 to 2014, using the Driver-Pressure-State-Impact-Response (DPSIR) framework [25]. To achieve this objective, we addressed two specific questions: (1) what are the LULCC patterns across geographic and biophysical settings through time, emphasizing in trends on native ecosystems as sentinel habitats, specifically how landscapes are being transformed over time, in terms of the rate, magnitude and direction of those changes and (2) what is the combination of environmental and anthropic factors that better explain the different landscape transitions.

## Materials and methods

### Study area

Pedro Moncayo county is located in the western branch of the Andes in northern Ecuador (Fig 1). This county is characterized by a wide elevation gradient (2400-4400 m a.s.l) and a management regime that varies in intensity depending on the elevation [26]. The higher altitudinal zone (above 3300 m) is dominated by native ecosystems, represented by paramo and highland montane forests [27]. The middle altitudinal area (2800 −3300) has been intensively used for agriculture and livestock through time, causing severe soil degradation [27,28] and the lower lands are characterized by a shrub dominated dry ecosystems (Figs 1 and 2).

**Fig. 1.**
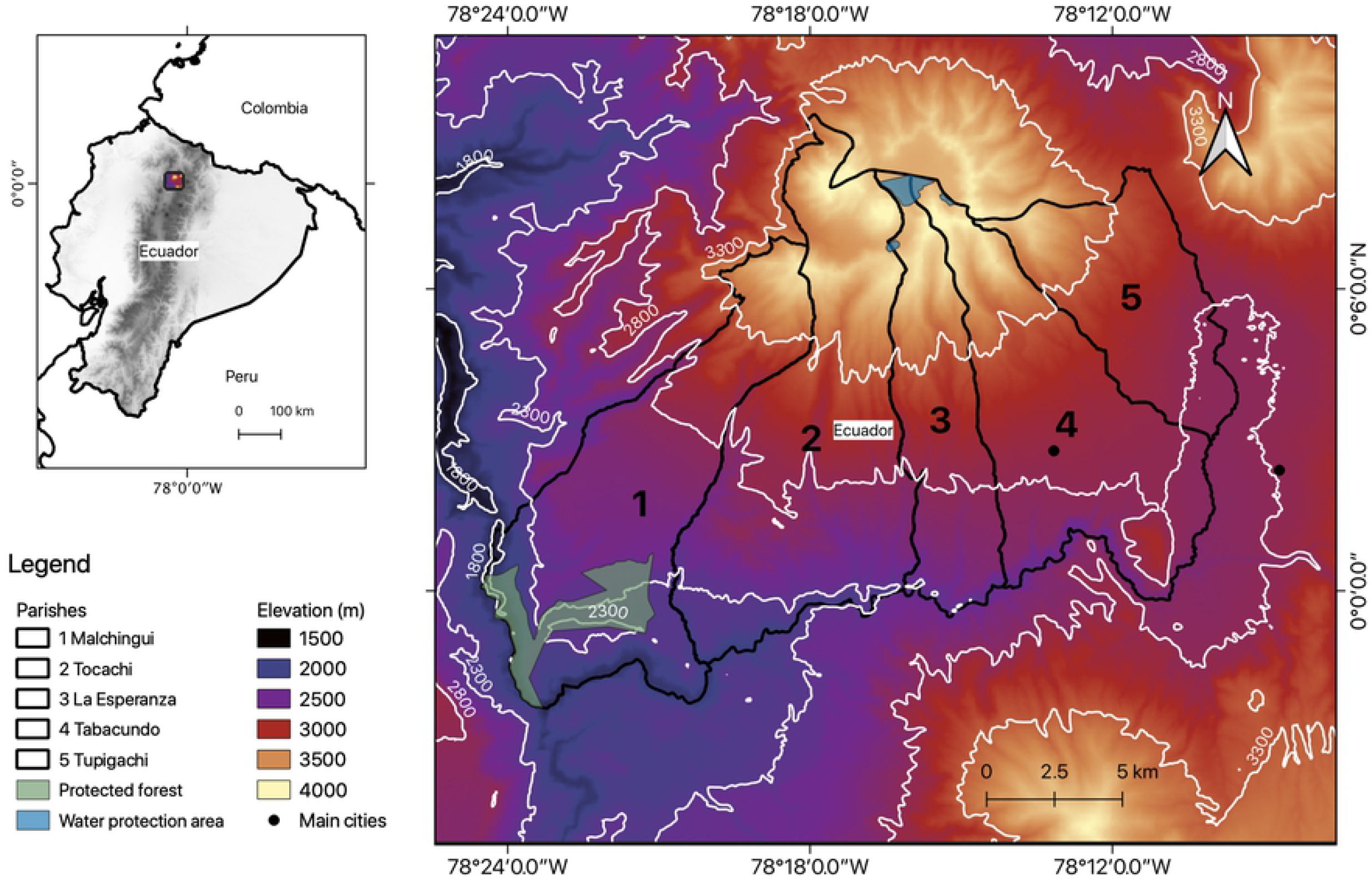
Location of study area (Pedro Moncayo county).

**Fig 2.**
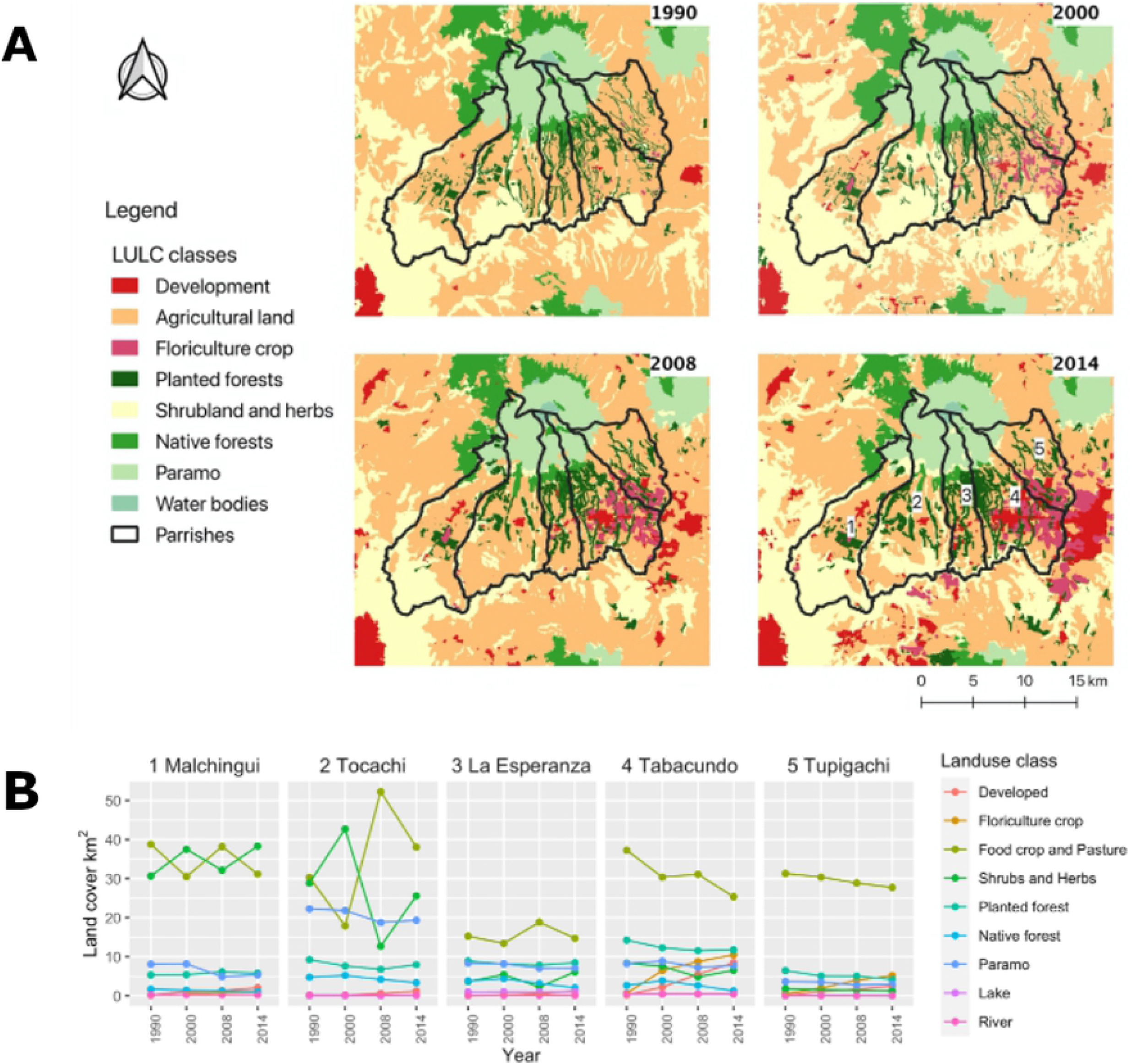
A. LULC maps of Pedro Moncayo county thoughout the periods of study (1990, 2000, 2008 and 2014). B. Land extent changes through time in the Pedro Moncayo county by administrative zones at the parish scale.

The studied territory has a total surface area of 339 km^2^, which is divided into 5 parishes that have an east to west geographic arrangement, depicting the same elevation belts previously described (Fig 1), however show different levels of production development and population trends; parishes located to the west portray a local economy based on subsistence agriculture and lower population growth, whereas the eastern parishes are attracting a growing population, have a more concentrated urban development, more irrigation systems and it harbors an expanding agro-industrial sector [26].

Pedro Moncayo is characterised by a typical climate of the tropical Andean region, with low annual variability but significant changes between night and day [29], in addition because of its elevation gradient, quarterly midday maximum temperatures could range from 14°C to 24°C and minimum nighttime temperatures could range from 4°C to 17°C [29]. In contrast, the precipitation pattern follows a bimodal peak of heavy rains concentrated from October to November and April to May, followed by a dry period of low precipitation from June to September; quarterly precipitation could range from 0 mm to 225 mm, and depending on the season, the extension of the territory could shift to a different hydrologic regime [29]. For instance, from April to June the majority of the territory could have more than 200 mm of precipitation, whereas in the quarter of July to September most of the area receives less than 75 mm of precipitation [29].

Approximately 4 % of the county’s territory is under conservation or environmental management, including the Jerusalem Protected Forest which occupies 1110 hectares of dry ecosystems in the county’s lowlands, and the Mojanda Lacustric complex, protecting only 26 hectares of highland ecosystems and water sources (Fig 1) [30].

Although, at present the majority (58.1%) of the territory of Pedro Moncayo is dedicated to traditional agricultural activities mainly for growing cereals, maize and potatoes, the economy of the region is based on the production and export of flowers (mainly roses) using greenhouse infrastructures [30]; small and medium-scale agriculture and livestock ranching are lower in terms of labor absorption, technology incorporation and productivity [27].

### LULC datasets

Official Land Use Land Cover (LULC) maps from the Ministry of Environment of Ecuador (MAE) of four periods of time: 1990, 2000, 2008 and 2014 were used to generate polygon vector from Landsat (TM) images with a spatial resolution of 30 meters and a temporal resolution of 16 days [31]. The LULC official classification encompasses a 2-level hierarchical scheme, based on the IPPC classes in combination with a taxonomy agreed by the entities in charge of generating land cover information in Ecuador [31]. Despite possible drawbacks to the LULC datasets, such as the existence of classification errors and uncertainties [32], its accessibility and availability at different time spans offer considerable advantages for studying land cover changes [33].

Five typologies not well represented in the official LULC dataset for the study region were digitised to improve map accuracy [34]. These included: planted forests, developed areas (populated zones), horticulture (areas represented by greenhouses) and natural water bodies. Composite LANDSAT images from our study area were obtained from the periods of interest (1990, 2000, 2008 and 2014) using the Code Editor of Google Earth Engine [35]. Results of the digitation process were overlapped over the LULC official vector layers from the periods of interest and further rasterized. Digitation, rasterization and overlapping analysis were conducting in QGIS 3.10 [36].

For our LULC change analysis we used a modified categorization from MAE-MAGAP [31], we combined level 1 and 2 official LULC taxonomy (S1Table). Briefly we aggregated all the agricultural level 2 typologies into agricultural land, and as suggested by MAE-MAGAP [31], we included pasture to this LULC class since in the highlands of Ecuador there is a system of rotation from pasture to agricultural fields along the cropping cycles. In addition, because the study area corresponds to the major center of floriculture production for the export market in Ecuador [37,38], we added floriculture crop, as a separate typology from the agricultural land. As a result, the identified LULC classes were 1) developed, 2) floriculture crop, 3) agricultural land, 4) planted forest, 5) shrubland and herbs, 6) native forest, 7) paramo and 8) water bodies (S1 Table).

### Land Use and Cover changes

Firstly, we mapped and estimated the land area occupied by each LULC class through time and the percentage change (C %) in each land-use class was calculated by dividing the area difference between the latest and the base year of each class by the coverage area in the base year and multiplying by 100 [24].

Then, LULC changes were estimated for three periods of analysis: for 1990-2000 (T1), 2000-2008 (T2) and 2008-2014 (T3). Furthermore, to analyze the succession of LULC classes in these periods of analysis, we used discrete-time, finite-state, homogeneous (stationary) Markov chain models, which have been widely used to model LULC changes [39-42]. The Markov Chain probability Matrix was estimated, using the markovchain R-package [43] for five administrative zones (at the parish level) and across four elevation bands. By applying a Markov chain model for three periods of analysis to land use classes it is possible to observe conversions between them when values are higher than 0.5); in contrast the stability probability is observed when higher values are compared between the same LULC class, representing the probability of remaining in the same class in the consequent time period, given the present state of the class. The spatial patterns of LULC change across administrative zones were obtained from an overlay procedure of the LULC maps with the polygons of parishes from the studied Pedro Moncayo county, which were downloaded from the official reference [44]. In the same way, to understand the patterns of LULC change across elevation classes, first the National Digital Elevation Model at a 30 m spatial resolution [45] was clipped to the study area, after that, the resulting image was further reclassified according to elevation bands, with an interval of 500 m [46,47], resulting in the following four elevation bands <2300, 2300-2800, 2800-3300, >3300 (Fig 1); these groupings take into consideration the mean medium of relief surface roughness, type of forests and the presence of urban areas [46]. Finally, the LULC classification for each year was layered over both (1) the reclassified elevation map, and the (2) reclassified administrative map. Spatial data assimilation, processing and overlaying analysis were conducted in the R environment [48].

### Drivers of change

To understand what predictors could explain the LULC dynamics we tested a set of five group of variables that have previously been reported as possible drivers of change [7,12] and were described in the conceptual framework that interconnects: Driving forces – Pressure – State – Benefits / Impacts-Response adopted by the European Environment Agency [25]; a scheme also used by the Ministry of the Environment of Ecuador as a tool to guide the formulation and adjustment of policies to foster biodiversity conservation in Ecuador [49]. Within this approach, we compiled a dataset of 13 variables of LULCC ranging from (1) socio-economic, (2) topography, (3) anthropic pressures to natural ecosystems, (4), climate and (6) governance decisions toward landscape development (Table 1).

**Table 1.**
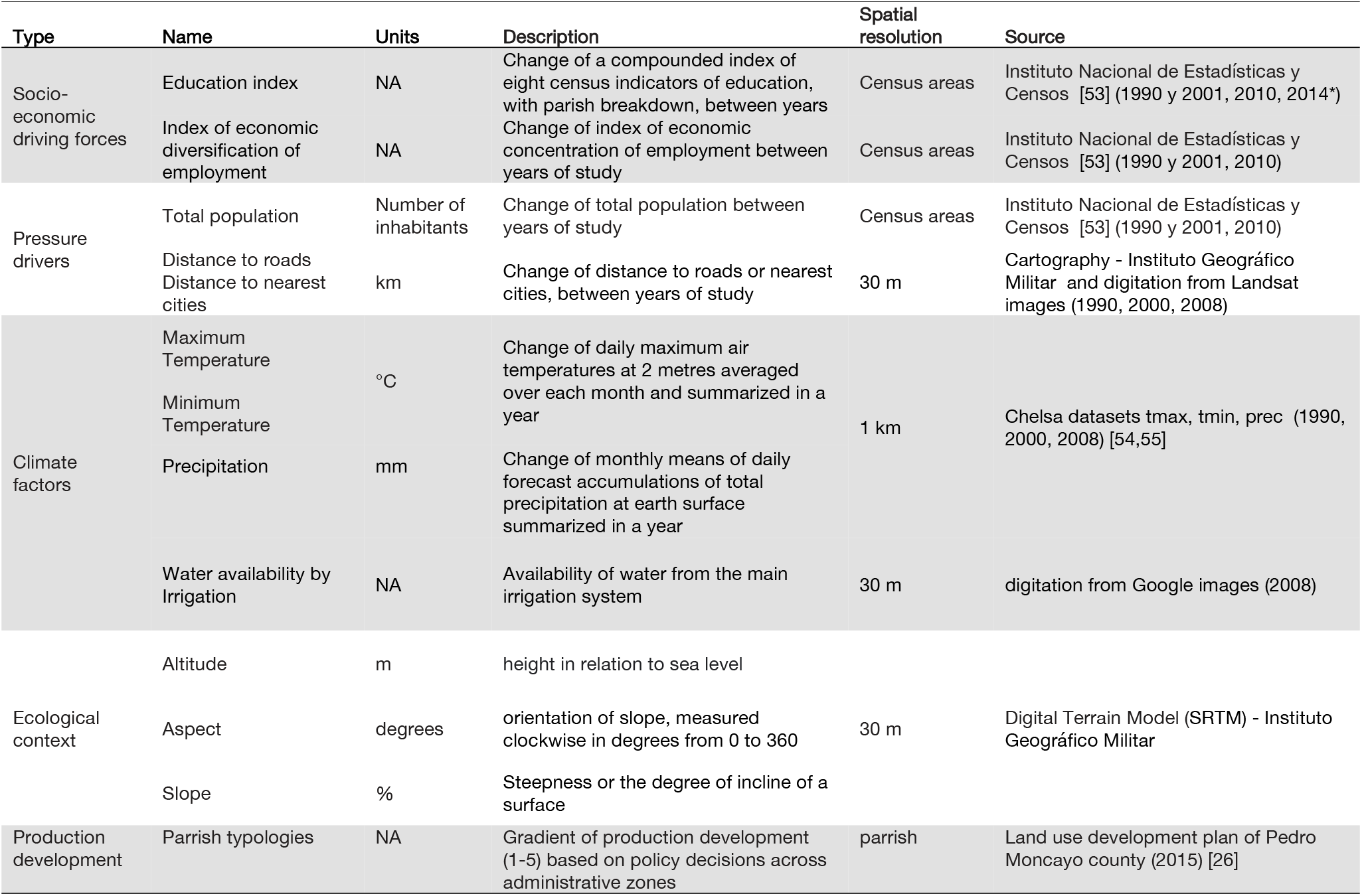
Predictors included in the General Additive Model to explain LULC probability of change.

In order to increase the number of units of analysis within parishes, all these variables were obtained at the spatial resolution of census area [50]. After all the spatial data assimilation, processing and visualisation necessary to obtain the drivers at the spatial unit of analysis, we carried out a reduction dimension procedure using Principal Component Analysis [51] of the drivers of change to summarize the distinct variables within the grouping drivers, all these procedures were completed using R software (version 3.2.3) [48]. Correlated variables were screened for the total variation explained by the first principal axes, and used to remove correlated variables [52]. Coordinates of the principal components that accounted for more than 60% of the variation were then used as explanatory variables in a subsequent statistical model.

### Statistical analysis

We synthesized and incorporated the different grouping drivers into a statistical model to improve LULC predictions and inform decision making by carrying out multivariate analysis using Generalized Additive Models (GAM). GAMs are an approach used extensively in environmental modelling and provide great scope to model complex relationships between covariates [56,57]. We used GAM regressions to elucidate two types of transitions in our study area: 1) the probability of natural ecosystems loss, and 2) the probability to change towards anthropic environments. The LULC trends evaluated as response variables within the first approach were the probability of loss of native forest, paramo and shrubs and herbs; complementary, the second approach tried to explain what drivers could cause the transitions towards developed areas, floricultural crops and food crops pastures. We did not include transitions toward planted forests because this LULC demonstrated to be very stable during the periods of analysis. As explained in the previous section, the explanatory variables for each GAM were the coordinates of the PCA that explained more of the 60% of the variation in the multivariate matrix.

The computational methods for the GAM modelling were implemented from the cran repository ‘mgcv’ package [58], since in our study the response variable is a probability ranging from 0 to 1, we used the GAM family as a Beta Regression, as suggested by this type of data [59]. For the smoothing basis function, we use the penalized cubic regression spline to lower computation cost and avoid overfitting; the smoothing parameter estimation was restricted maximum likelihood (‘REML’), typically used for smooth components viewed as random effects [56]. After checking the results of different models using distinct methods for selecting the number of knots (default, cross validation and manual adjustments), we selected the more conservative models, setting the number of knots in three to be flexible enough to allow the models to fit simple curve relationships and preventing spline curves with complex overfitting estimates, which would have limited the interpretation ability from the ecological perspective. We presented the results of the GAMs with Partial Dependence Plots (PDP) using the ‘mgcv’ R-package [58] to determine which variable best explained the variation in LULC change [56].

## Results

### Coverage area for each year

Agricultural land was the most representative LULC type in the study area, followed by shrubs and herbs (Fig 2). Both LULC types were very dynamic over the different periods of analysis, agricultural land ranged from 35 to 50% of the total area, depending on the period of analysis (Table 2), and shrubs and herbs varied from 16 to 28% of the total area, depending on the period analyzed (Table 2).

**Table 2.** Changes in land cover classification in Pedro Moncayo county from 1990 to 2014

| LULC<br>TYPE | YEAR | 1990 |  | 2000 |  | 2008 |  | 2014 |  | 2014-1990<br>% |
| --- | --- | --- | --- | --- | --- | --- | --- | --- | --- | --- |
|  |  | km<sup>2</sup> | % | km<sup>2</sup> | % | km<sup>2</sup> | % | km<sup>2</sup> | % | Change |
| Developed |  | 0.58 | 0.17 | 4.72 | 1.39 | 9.61 | 2.84 | 15.54 | 4.60 | 2569.55 |
| Floriculture crop |  | 1.19 | 0.35 | 9.44 | 2.79 | 14.06 | 4.16 | 16.75 | 4.95 | 1305.89 |
| Food crop and<br>pasture |  | 152.92 | 45.20 | 122.58 | 36.23 | 169.29 | 50.04 | 137.03 | 40.51 | -10.39 |
| Planted forest |  | 44.16 | 13.05 | 38.56 | 11.40 | 37.35 | 11.04 | 38.24 | 11.30 | -13.42 |
| Shrubs and Herbs |  | 73.33 | 21.68 | 94.74 | 28.00 | 53.40 | 15.78 | 77.75 | 22.98 | 6.02 |
| Native forest |  | 12.96 | 3.83 | 15.11 | 4.47 | 11.29 | 3.34 | 7.77 | 2.30 | -40.03 |
| Paramo |  | 50.62 | 14.96 | 50.60 | 14.96 | 40.73 | 12.04 | 42.51 | 12.57 | -16.03 |
| Lake |  | 1.48 | 0.44 | 1.52 | 0.45 | 1.52 | 0.45 | 1.66 | 0.49 | 12.29 |
| River |  | 1.04 | 0.31 | 1.06 | 0.31 | 1.04 | 0.31 | 1.04 | 0.31 | 0.09 |
| Total |  | 338.29 | 100.00 | 338.33 | 100.00 | 338.29 | 100.00 | 338.29 | 100 | 0.00 |

**Table 2.** Summary of the results of the General Additive Models to elucidate drivers of change for the six LULC transition models in Pedro Moncayo county.

| Drivers | native forest loss |  |  | paramo loss |  |  | shrub loss |  |  | urbanization |  |  | Floriculture expansion |  |  | Agriculture expansion |  |  |
| --- | --- | --- | --- | --- | --- | --- | --- | --- | --- | --- | --- | --- | --- | --- | --- | --- | --- | --- |
|  | p value | chi sq | edf | p value | chi sq | edf | p value | chi sq | edf | p value | chi sq | edf | p value | chi sq | edf | p value | chi sq | edf |
| socioeconomic PC1 | 0.000 | 19.65 | 1.8 | 0.721 | 0.00 | < 1 | 1.000 | 0.00 | < 1 | 0.468 | 0.00 | < 1 | 0.307 | 0.05 | < 1 | 0.812 | 0.00 | < 1 |
| ecological context PC1 | 0.000 | 13.32 | 1.4 | 0.000 | 23.89 | 1.5 | 0.426 | 0.00 | < 1 | 0.000 | 16.69 | 1.7 | 0.001 | 9.62 | 1.4 | 0.000 | 18.48 | 1.9 |
| climate factors PC1 | 0.889 | 0.00 | 1.0 | 1.000 | 0.00 | < 1 | 1.000 | 0.00 | < 1 | 1.000 | 0.00 | < 1 | 0.774 | 0.00 | < 1 | 0.814 | 0.00 | < 1 |
| climate factors PC2 | 0.336 | 0.00 | < 1 | 0.791 | 0.00 | < 1 | 0.374 | 0.00 | < 1 | 0.609 | 0.00 | < 1 | 0.018 | 4.23 | < 1 | 0.039 | 3.06 | < 1 |
| pressure drivers PC1 | 0.547 | 0.00 | < 1 | 1.000 | 0.00 | < 1 | 0.000 | 17.02 | 1.76 | 0.000 | 26.71 | 1.8 | 0.936 | 0.00 | < 1 | 0.000 | 15.21 | 1.3 |
| pressure drivers PC2 | 0.189 | 0.721 | < 1 | 1.000 | 0.00 | < 1 | 0.364 | 0.00 | < 1 | 0.048 | 0.00 | 1.2 | 0.144 | 1.24 | < 1 | 1.000 | 0.00 | < 1 |
| parrish gov | 0.507 | 0 | < 1 | 0.118 | 1.46 | < 1 | 0.000 | 25.53 | 1.53 | 0.386 |  | < 1 | 0.000 | 10.41 | 1.2 | 0.000 | 31.00 | 1.8 |
| Deviance explained | 21.00% |  |  | 20.80% |  |  | 39.90% |  |  | 36.50% |  |  | 25.00% |  |  | 41.80% |  |  |
| R-sq.(adj) | 0.22 |  |  | 0.15 |  |  | 0.29 |  |  | 0.30 |  |  | 0.21 |  |  | 0.33 |  |  |

Overall natural ecosystems, which are mainly represented by native forests and paramos, decreased from 1990 to 2014 (Table 2), there was a 40% and 16% decrease of native forest and paramo cover when compared the first and last periods of study (Table 2); but, by the last period of study, areas of paramo still represent an important part (13%) of the study territory. Natural water bodies (lakes and rivers) showed high persistence over time (Table 2).

Developed areas and floricultural crops were continuously increasing over time, and although they were poorly represented in the first period of analysis (less than 0.4% in 1990), by 2014 they represented almost 5% of the study area (Table 2), demonstrating a 26 and 13 times fold of increase from 1990 to 2014, respectively.

Landscape dynamics through time was not homogenous along the study area, instead it shows a geographic pattern (Fig 2). Expansion of developed areas and floriculture crops occurred mainly in the southeastern part of the studied region (Fig 2). The greatest degree of loss in native forests and paramos occurred in the northeast (Figs 2 and 3), where there is almost no paramo left due to the expansion of agricultural land.

**Fig 3.**
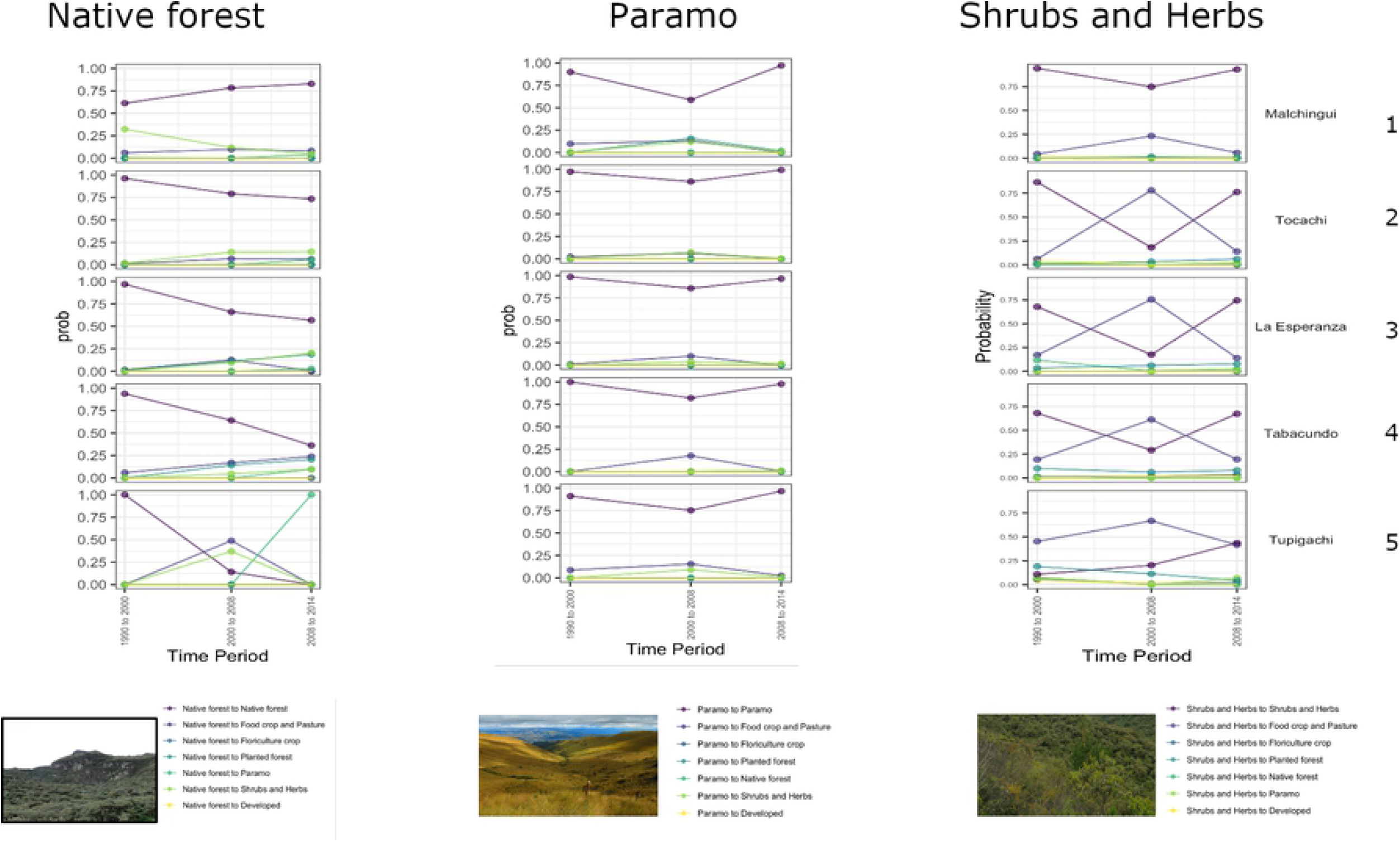
Transition probability of native ecosystems through time in Pedro Moncayo county, at the parish level

### Land-change dynamics through time

#### Transitions of native ecosystems

In general, as expected the stability of native forests is decreasing through time in the entire territory (Fig 3), with the exception of the western parish where the probability of remaining in this LULC class increases through time probably due to agricultural land abandonment (Fig 3). In contrast, areas located in the east tend to have lower values of stability through time and higher probabilities to change to paramo and agricultural land; this pattern was more evident in the last period evaluated (2008-2014) (Fig 4). Additionally, this trend is more evident along elevation bands; where native forests located above 3300 m showed a lower probability of remaining as forest along the years (Fig 4) and in the 2800-3300 altitudinal belt there is a high probability of converting native to planted forests, especially in the center of the territory (Fig 4).

**Fig 4.**
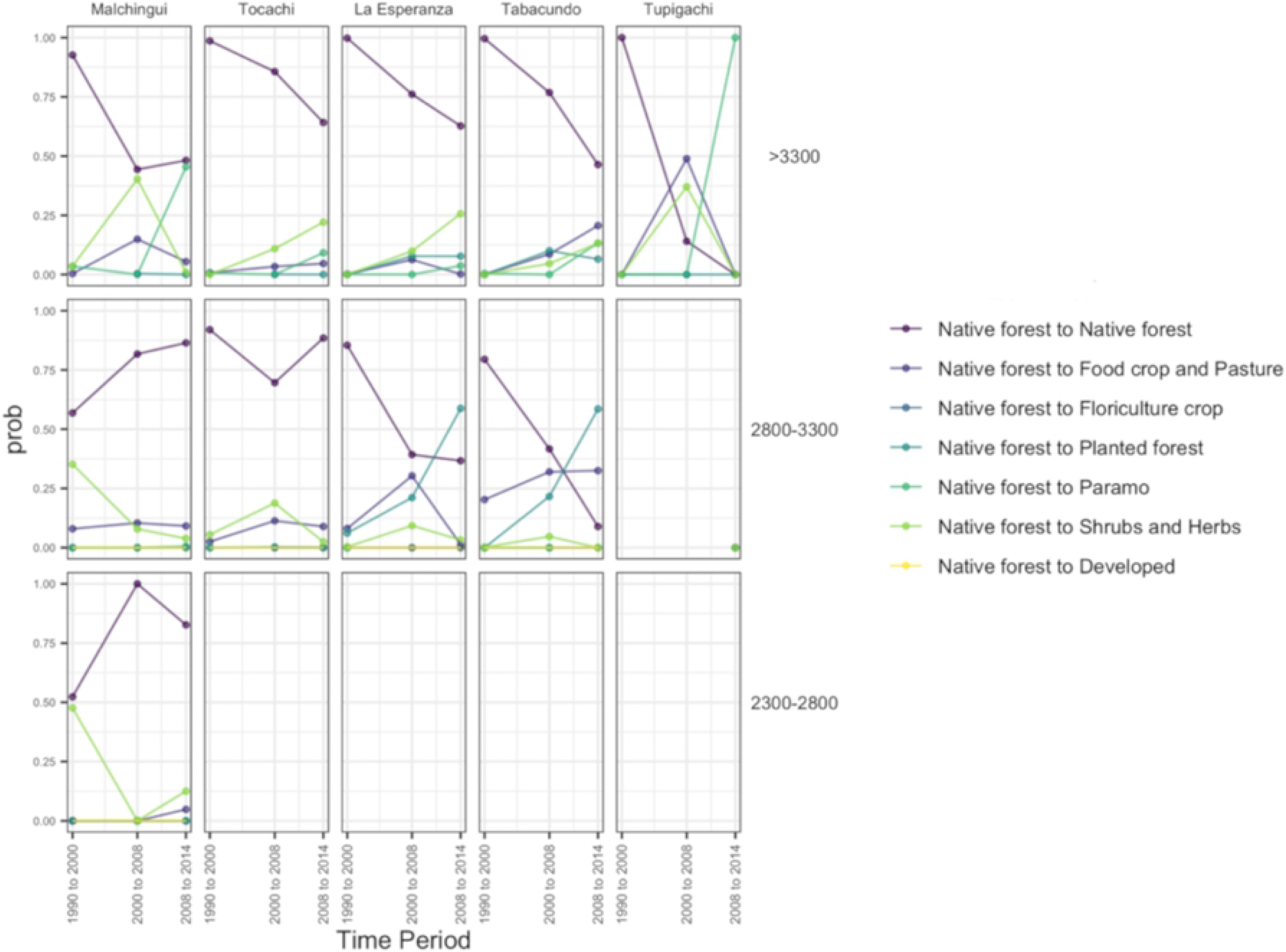
Transition probability of native forests through time in Pedro Moncayo county, by altitudinal bands at the parish level.

Furthermore, shrubs and herbs show variable change throughout the study period (Fig 3). In the majority of administrative areas, the stability of shrubs and herbs decreased (from values around 0.75 to values close to of 0.25) in the second period of evaluation (2000-2008) and increased again in the last period (2008-2014). Across all elevation belts, this LULC class tend to follow a dynamic trend changing back and forth with the agricultural land; however, this pattern was not observed in the eastern parish at the elevation belt of 2800 – 3300, where the landscape seems to have a high probability of remaining as agricultural land (S1 Fig).

In contrast, paramo is the most stable among all natural ecosystems evaluated, although a slight decrease in stability was observed from values above 0.90 to around 0.75 in the second period of analysis (2000-2008) (Fig 3), the probability of remaining in the same land use class increased by the last period of analysis (2008-2014). Since this ecosystem is characteristic of highlands (above 3000 m) the transition probabilities were only observed for the two higher elevation belts evaluated and their stability seems to be increasing in the administrative zone located in the western part of the territory (S2 Fig).

#### Transitions to anthropic environments

Developed areas demonstrate a differential trend through time in the study area (Fig 5). In the western areas of the territory (Fig 5) the stability of this LULC class decreased in the second period of evaluation (2000-2008) and significantly increased again in the last time period (2008-2014), in contrast, the parishes located to the east exhibit a more stable probability of remaining as developed areas through time probably due to the proximity of the larger towns (Fig 5). Since the territory studied is in general a rural area, there is a dynamic trend towards converting agricultural to urban areas, which follow a geographic pattern (Fig 5).

**Fig 5.**
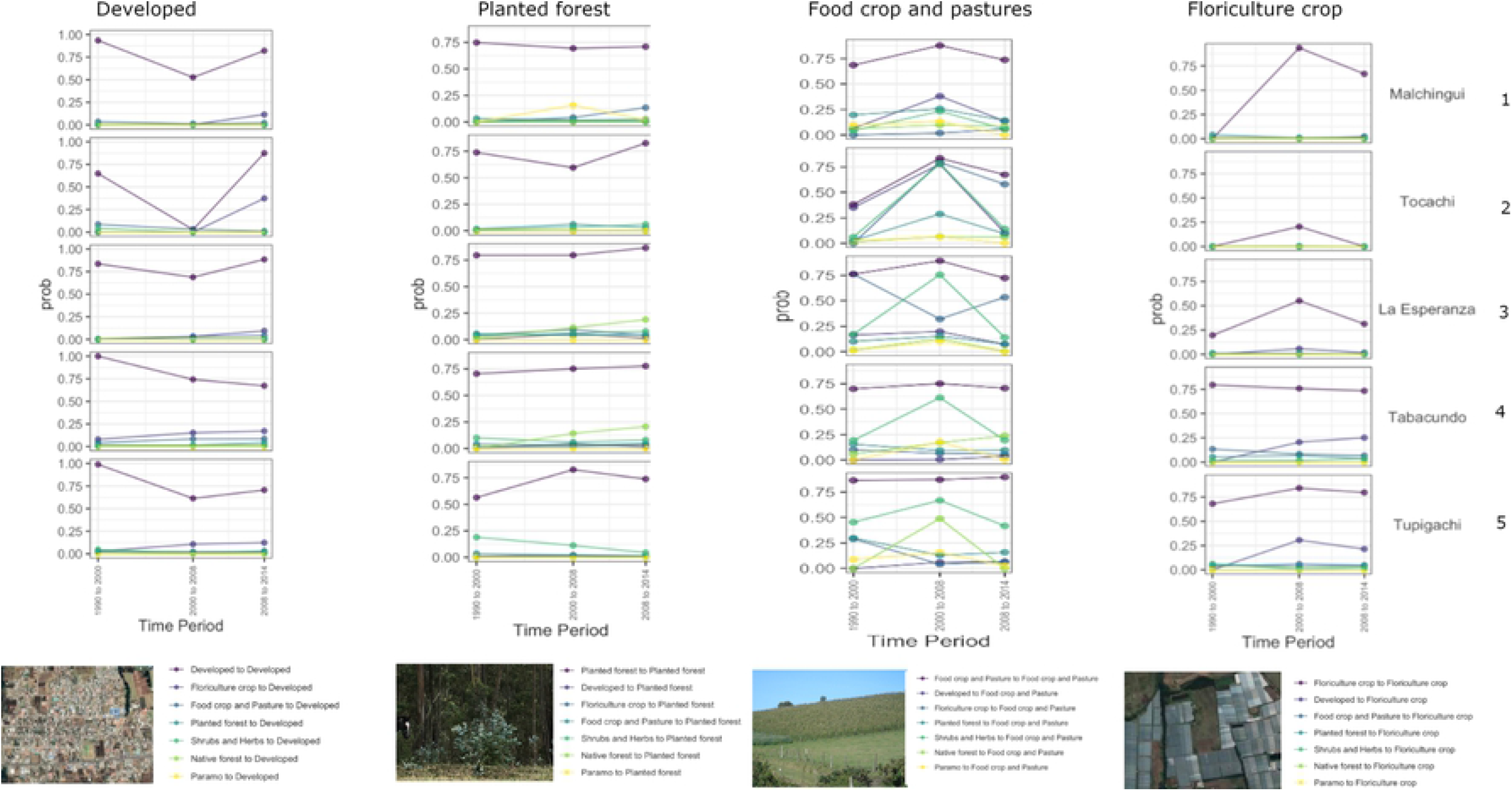
Transition probability to anthropic environmets through time in Pedro Moncayo county, at the parish level

To the east and center of the study area, floriculture crops have not been fully established because it tends to change toward Agricultural land (Fig 5); in contrast this LULC type located in the eastern parishes is more stable with values around 0.75 throughout the period of study (Fig 5).

Agricultural land is a very stable land use class throughout the study period in the administrative zones located in the center and eastern part of the study area with values ranging above 0.77 (Fig 5); the stability of this land use class in the west followed a dynamic trend through time: in the first period of analysis (1990-2000) it was lower than in the second period of analysis and it increased again by the last period of analysis. In contrast, planted forests depict a very stable land use trend thorough time across the territory, their probability of remaining in the same land use class ranges from 0.6 to 0.90 (Fig 5).

### Drivers of change

The drivers of change, which are mainly the result of a dimension reduction of distinct variables organized in grouping drivers by PCA analysis, displayed different spatial distribution within the study area (S3 and S4 Figs), depicting a territory with contrasting patterns. The details of the spatio-temporal distribution of the drivers of change are presented in S3 and S4 Figs.

Table 2 describes the results of the different LULC transitions studied and their main explanatory variable; the General Additive Models demonstrated different results when explaining each LULC transition (Table 2). The lowest total variance explained (21.00%) corresponded to the native forest loss model and the largest value (41.80%) was for the agricultural expansion model. Overall, the most relevant parameters explaining LUCC in the region were the ecological context grouping driver (which incorporates elevation and slope), this grouping driver was highly significant for the majority of the transitions studied (p < 0.001, Table 2), with the exception of the Shrub and Herbs expansion. In contrast, the climate grouping driver PC1 (which depicts mostly the variation of precipitation and minimum temperature) was not significant in any model (p >0.05).

For the native forest loss model, the most important grouping drivers (p < 0.001) were the socioeconomic and the ecological context grouping drivers (Fig 6, Table 2). For instance, paramo loss was only explained by the variation in elevation and slope (ecological context PC1) (Table 2, S5 Fig). Figure 6 shows the GAM PDPs for the native forest loss model and indicates that the probability of native forest loss increases as land aspect PC1 increases, in other words, when elevation and slope increases. In contrast, when the socioeconomic variables have low and high values the probability of forest loss increases, although the confidence interval for lower values in the socioeconomic drivers has higher values.

**Fig 6.**
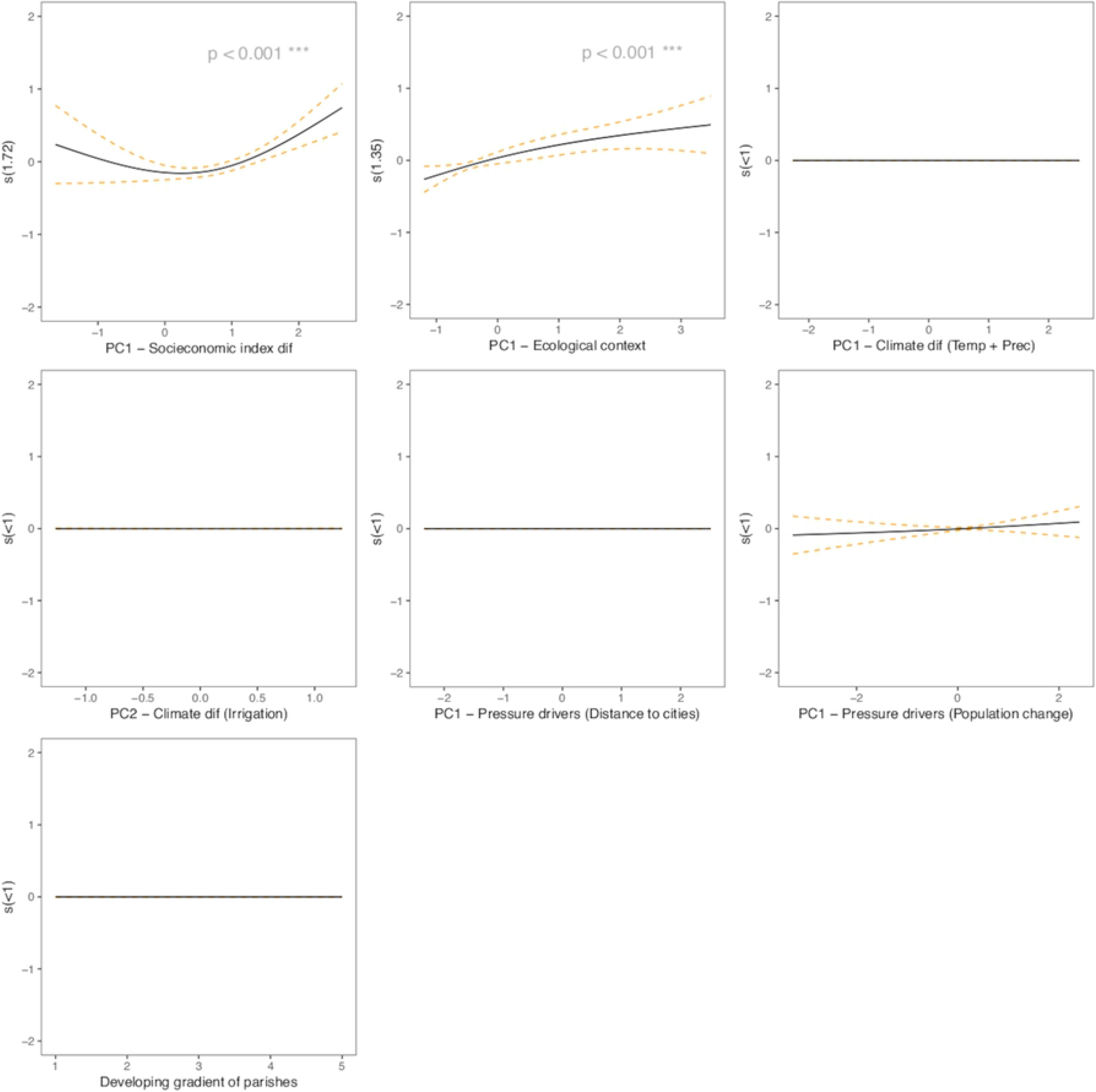
Generalized additive model partial dependence plots for forest native loss. Each plot shows a covariate and their partial dependence on probability of native forest loss in the context of the model. The y axis shows the mean of the probability of native forest loss and the x axis the covariate interval. The gray area represents the 95% confidence interval.

Some transition models were explained by similar grouping drivers such as: shrub loss and agricultural expansion (Table 2), which also show a contrasting pattern in their response variables, in such a way that when was prevalent an increase in agricultural areas it was a decrease in shrub and herb extension (Fig 2); these models depicted the following grouping drivers as significant parameters (p<0.001): pressure drivers (PC1) and a variable that describes differences development of the distinct administrative areas within the study area (Table 2). S7 and S8 Figs shows the GAM PDPs for shrub and herb loss and agricultural expansion respectively, and they reveal that high probabilities for these transitions are related to medium values of elevation and slope (ecological context drivers), additionally as the main local cities are further away increases the probability to convert natural areas to agricultural land, and there is a linear increase in rates of change to agricultural land with the gradient of development at the parish level.

Variables leading to the highest change in the probability of transition to floriculture crops comprise the ecological context grouping driver, the climate PC2, which includes water irrigation and the development gradient across parishes. Floriculture crops increase as elevation and slopes decrease. Complimentary, when more water is available through irrigation, the probability of establishing floriculture crops increase (S9 Fig).

For instance, the urbanization transition model was explained by ecological context and pressure grouping drivers PC1 (p< 0.001) (Table 2). Urban transition probabilities decrease significantly (p<0.001) with altitude and slope, it also significantly decrease (p<0.001) with the distance to the city centers (pressure PC1), and with higher values of total population change (pressure - PC2) (S10 Fig).

## Discussion

Even though major milestones for landscape transformation occurred centuries ago in the Ecuadorian highlands [6,60–62], our study demonstrates that, at present, a combination of environmental variables and human induced factors still have an impact on an even LULC transformation as reported for other mountainous landscapes of Latin America [7,9,10,12]. The studied area depicts a rural Andean landscape dominated by an agricultural matrix which contains important areas of shrubland and paramo, accompanied by patches of remaining native forest, which is consistent with other current landscapes in the Tropical Andes [7,12,24,62].

We also found a geographic pattern of LULC transition across the study area; we report a clear deforestation pattern of native montane forests located below 3300 m.a.s.l, an unexpected high stability pattern of paramos for the majority of the studied territory and a dynamic transition between agricultural land and shrubland was found; in addition, we found an exponential increase in urban land and floriculture crops in the eastern part of the territory. This result is striking because of the small spatial scale where the changes occur; our study area encompasses only 334 km^2^ compared to other landscapes studied in Central Ecuador [62], in the Peruvian Puna [24] or in Colombia [12], where the land extent are 10, 120 and 800 times larger than our studied territory.

We estimated a paramo loss of 16% from 1990 to 2014 in Pedro Moncayo county (Table 1), this result is consistent with the findings of loss (13%) in a nearby territory [63]. Although most of the studied territory depicted a high stability pattern of paramo, as also described for a highland landscape of central Ecuador (Ross et al 2017), our results also demonstrated a hotspot of paramo conversion to agricultural land concentrated in the northeast. In contrast, our results are strikingly different with the land cover patterns observed in other paramos in the region, where a more widespread agricultural use of paramo was observed [64-66]. Although, another common transition reported for paramos in the Ecuadorian mountains is to exotic timber plantations [18], this trend was not apparent for our studied territory.

We found a 40% montane forest loss from 1990 to 2014, and the Markov chain model demonstrated a very low probability of persistence of this ecosystem in the majority of Pedro Moncayo county (Figs 2 and 4); this is consistent with the general trend of deforestation and degradation of mountain forests in the Tropical Andes mainly explained by agricultural expansion [67]. We also found that the highest chances of loss occur in the altitudinal band of 2800 to 3300 (Fig 3); these findings are in accordance with the findings described for other representative highlands in central Ecuador [62]; however, LULC change studies carried out in more isolated landscapes of central and southern Ecuador reported deforestation hotspot for lowland montane forests and afforestation transition in upper altitudinal areas [10,68]; additionally, higher rates of deforestation was also observed in lowland forest of Colombia, in the Napo region along the Ecuadorean border [12].

Mountain forests are considered one of the most threatened forests in the tropics [69,70], which are also highlighted as a global priority for conservation due to their great biodiversity and high level of endemism [71], and its vital role in the provisioning of different ecosystem services in the region [72,73]. However, if the trends demonstrated by the Markov model are maintained for this territory, there is a high probability that the remnant montane forests will be permanently lost in a few years, posing a greater threat to the already vulnerable biodiversity [69] and limiting the capacity of these ecosystems to provide services in the county, such as provisioning and regulating of freshwater, “wild foods” and many other non-timber forest products [22,74], as described for other latitudes [75–77].

Along with this deforestation trend, we observed a dynamic and opposite transition between agriculture areas and shrubland, this pattern was more evident for the parishes located in the center of Pedro Moncayo county and along the elevation bands between 2300 to 3300 m.a.s.l. This pattern could demonstrate a gain of secondary vegetation, probably due to a temporal abandonment of agricultural areas, follow by a net gain of agricultural land which has been observed in other Andean systems of Colombia [12] and Central America [78].

We found that urban areas are dramatically increasing in the eastern part of the territory (Fig 1, Table 1); we reported a 25-fold increase in urban cover from 1990 to 2014. This pattern is following the global trend of urban expansion [79], but the rate of extension is even faster than what has been reported for many cities around the world [80] and in small urban centers [81] [79]. In our study, higher probabilities of urban land expansion were explained by increases in population, proximity to urban centers and occurred at lower elevations and slopes in previous crop land. This pattern has been observed in other regions of South America, where urban expansion is taking place largely on agricultural land [79], a zone characterized by areas of lower altitude and slope, which in the Andean zones correspond to the more fertile valleys between mountains.

Another interesting finding was the exponential expansion of flower cultivation cover reported for Pedro Moncayo county (Table 1, Fig 2). We described a 13-fold increase in total land area of greenhouse floriculture from 1990 to 2014 (Table 1); this expansion was observed primarily in the eastern parishes of the territory (Figs 1-3), which are located contiguous to Cayambe county, another center for the development of this activity in Ecuador [63]. This region, situated in Pichincha Province, has an equatorial location and has optimal sunlight conditions (long hours of daylight) and an ideal highland climate (abundant sunshine, warm days and cool nights) which make it possible to produce some of the highest quality flowers in the world [37,38].

Our analysis suggests that in addition to the ecological context variables, another driver that explain the floriculture expansion pattern is the water availability by irrigation, depicted by the geographic pattern of irrigation in the lower eastern part of the studied territory, creating a better environment for growing crops which would have been limited by natural precipitation as demonstrated by [82] to increase yield in many crops. This irrigation canal transports water from the glacier of a snow-capped mountain located in a contiguous territory, corresponding to the neighboring county (Cayambe), and reaches only to the center of the territory and distributes water to lower elevations, therefore only the area situated to south-east receives water for irrigation.

We found that ecological context variables (elevation and slope) are the most important drivers for all LULC transitions. For instance, native ecosystems transitions (including the models to explain loss of native forest and paramo) and agricultural expansion were both significantly related to changes in elevation and slope, in such a way that the probability of native ecosystems loss and the probability of agriculture expansion increase with elevation and slope, until they reach a certain value where they level off (native forest and paramo models) and even decrease (shrubs and herbs loss and agricultural expansion models). These complementary trends suggest that the major pressure for native ecosystems in this highland system of northern Ecuador is still the expansion upwards of the agricultural-livestock frontier as also found in other Andean landscapes [12,62]. In addition, the expansion of urban areas and floriculture crops in the previous agricultural land, located at lower elevations of the eastern part of the territory could also make pressure for an expansion of the agricultural frontier in highland areas. Even though, we did not find evidence that climatic variation explained the LULC transitions, the effect of climate change could be stronger in a nearby future due to the extreme events predicted in the Tropical Andes [83], affecting the capacity of highland ecosystems to keep providing key goods and services to people [84].

The trend of native ecosystem loss associated to higher elevation and slope observed in this landscape of northern Ecuador could be attributed by its past patterns of land use as summarized by [3,85]; since the most striking transformation and loss of native ecosystems in Andean landscapes occurred centuries ago and it was even expanded in the mid-twenty century by the agrarian reform, current native ecosystems are only the remnant patches, localized at higher elevations and slopes [22]. However, the leveling off and the further decrease in the probability of native forest loss at higher values of topographic variables, could be explained by conservation measures adopted to restrict human activities in the upper mountain belt, like establishing protected areas [3,22,62] or implementing national or local policies to limit agricultural expansion [4] that have prevented the loss of high mountain ecosystems in other Andean regions [7,19].

Paramos and other high-elevation ecosystems (pristine native forest patches), which are ecosystems situated above 3500m in the northern highlands of Ecuador, are currently more valued by its importance for provisioning critical ecosystem services and, thus, in Ecuador have received special protection measures at the national [86,87] and local level [88].

Although, other studies [7,12,24] have also found that environmental variables such as topography were better predictors of woody vegetation change, arguing that these variables place physical limits on the types of land-use practices that are feasible in a region, the trends were different from those observed in our study in that these authors found that deforestation occurred in the lowlands, which are more appropriate for large-scale mechanized agriculture [7,12,24]. However, the dynamic transition trend between agricultural land and shrubland observed in our study, could be attributed to natural reforestation succession at high elevations (e.g., cooler temperatures, steeper slopes), which is consistent with other studies [7,68]. In our study, this pattern was also associated to variation in population change, which could be attributed to population migration dynamics within the territory; movements of farmers from higher mountainous zones to urban concentrated areas have been largely documented in different regions of Latin America and are the drivers associated to natural reforestation in higher elevations due to agricultural land abandonment [19]. This finding is consistent with the local demography dynamics, where urban population has tripled from 1990 to 2010 (from 3000 to 10000 inhabitants) while rural population has doubled (12000 to 23000 inhabitants) in the same period [26].

In many places where this landscape transition was reported it has facilitated ecosystem recovery in the highlands, likewise this has allowed maintaining the provision of ecosystem services for a growing urban population [19]. The dynamic conversion from agricultural land to shrubland in some highland areas of this landscape, explained by rural-urban migration, is consistent with the “Forest Transition Model” proposed by Mather [89], however in our study area the pattern is uneven, for instance native forests are decreasing in some areas, while shrubland is expanding in other areas, picturing a process of ecological succession before a fully recovered forest occurred. Maintaining and increasing native ecosystems in higher elevations and expanding urban and agriculture areas in the lowland and valleys raise new opportunities and challenges for conservation; however, the consequences of these spatial transitions have not been studied in depth [19].

We have considered a comprehensive set of factors characterizing landscape conversion dynamics, however some limitations concerning the scope of the drivers used for this analysis should be considered. The underlying driving forces affecting land-use transformations could also be attributed by production support policies geared toward internal market and exports [21,62], which were not included in our analysis. For instance, other studies suggest that the greenhouse floriculture expansion initiated in the 1990s was partly in response to favorable trade agreements but also due to diffusion of technologies from multiple sources and local entrepreneurship [38]. Flower cultivation is a land- and labor-intensive activity with high land productivity (that is, high market value of output per hectare) [37]. However, the gains in income have surely been offset by growing health and environmental problems posed by the intensive use of pesticides in flower cultivation [37].

All indications suggest that flower exports will continue to play a major and probably increasing role in Ecuador’s economy [37]; in fact, it is steadily expanding and it is causing land use changes in the territory; for instance, former important and traditional lands dedicated to livestock and food crop production, located in areas with aptitude to agricultural production and with access to irrigation systems have been transformed into green houses for flower cultivation, posing a the trade-offs between agricultural production and environmental concerns, including the asserted need for global land use expansion, and the issues of rural livelihoods and food security [30].

The assessment of local and regional patterns of current land use and past land cover conversion is the first step in developing sound land management plans that could prevent broad scale, irreversible ecosystem degradation. This characterization of landscape patterns through time and the analysis of their proximate drivers of landscape change enhance our understanding of how landscape pattern might influence ecosystem services [90] and point out that research and landscape management, zonation and ecological recovery/ restoration should become better integrated into land-use policy and conservation agendas at the local level to balance the multiple needs and benefits from Ecosystems of a growing population in a rural landscape of northern Ecuador [91].

## Acknowledgments

Thanks to the staff of the National Institute of Statistics and Census for sending the census cartography databases not available on the web page and for their advice in interpreting the coding of the databases. Thanks to Phoebe Lehmann Zarnetske for her sound suggestions on the available climate databases. And last but not least, thanks to the students of the School of Biological Sciences of the Central University who helped to collect information, especially Genesis Granja who directly supported in obtaining geospatial data related to the drivers of change.

## Author Contributions

Conceptualization: Paulina Guarderas, Franz Smith, Marc Dufrene.

Data curation: Paulina Guarderas.

Formal analysis: Paulina Guarderas, Franz Smith.

Funding acquisition: Paulina Guarderas, Marc Dufrene.

Investigation: Paulina Guarderas.

Methodology: Paulina Guarderas, Franz Smith.

Project administration: Paulina Guarderas.

Resources: Paulina Guarderas, Franz Smith.

Supervision: Marc Dufrene.

Validation: Paulina Guarderas, Franz Smith, Marc Dufrene.

Visualization: Paulina Guarderas, Franz Smith.

Writing – Paulina Guarderas.

Writing – review & editing: Paulina Guarderas, Franz Smith, Marc Dufrene.

## Supporting information

**S1 Figure. Transition probability of paramo through time in Pedro Moncayo county, by altitudinal bands at the parish level**

**S2 Fig. Transition probability of shrubs and herbs through time in Pedro Moncayo county, by altitudinal bands at the parish level.**

**S3 Fig. Spatial distribution of each grouping driver for the first period of analysis. Each map represents the PC1 from the Principal Component Analysis carried out for each grouping driver of change from the period 1 (1990 and 2000).**

**S4 Fig. Spatial distribution of each grouping driver for the second period of analysis.** Each map represents the PC1 from the Principal Component Analysis carried out for each grouping driver of change from the period 2 (2000 and 1990).

**S5 Fig. Generalized additive model partial dependence plots for forest paramo loss.** Each plot shows a covariate and their partial dependence on probability of native forest loss in the context of the model. The y axis shows the mean of the probability of native forest loss and the x axis the covariate interval. The gray area represents the 95% confidence interval.

**S6 Fig. Generalized additive model partial dependence plots for shrubland loss**. Each plot shows a covariate and their partial dependence on probability of native forest ln the context of the model. The y axis shows the mean of the probability of native forest loss and the x axis the covariate interval. The gray area represents the 95% confidence interval.

**S7 Fig. Generalized additive model partial dependence plots for agricultural transition**.

Each plot shows a covariate and their partial dependence on probability of native forest ln the context of the model. The y axis shows the mean of the probability of native forest loss and the x axis the covariate interval. The gray area represents the 95% confidence interval.

**S8 Fig. Generalized additive model partial dependence plots for floriculture transition**.

Each plot shows a covariate and their partial dependence on probability of native forest ln the context of the model. The y axis shows the mean of the probability of native forest loss and the x axis the covariate interval. The gray area represents the 95% confidence interval.

**S9 Fig. Generalized additive model partial dependence plots for urban transition**. Each plot shows a covariate and their partial dependence on probability of native forest ln the context of the model. The y axis shows the mean of the probability of native forest loss and the x axis the covariate interval. The gray area represents the 95% confidence interval.

**S1 Table. Land Use Land Cover classification scheme used to assess LULC change analysis** [31]

